# Passive acoustic monitoring of the rare orange-bellied parrot: automated detection using BirdNET-based transfer learning

**DOI:** 10.64898/2026.06.01.729442

**Authors:** Giselle Owens, Connor M. Wood, Thomas J. Hunt, Laura T. Bussolini, Annie Kreisl, Fernanda Alves, Dejan Stojanovic

**Affiliations:** Fenner School of Environment and Society, College of Systems and Society, Australian National University; Cornell K. Lisa Yang Center for Conservation Bioacoustics, Cornell Lab of Ornithology

**Keywords:** Orange-bellied parrot, *Neophema chrysogaster*, passive acoustic monitoring, BirdNET, automated bioacoustic classification, custom acoustic detector, wildlife monitoring, occupancy

## Abstract

Efficiently finding rare species is a perennial challenge in conservation science. The orange-bellied parrot *Neophema chrysogaster* is a rare mobile bird that is difficult to locate using traditional field survey techniques with human observers. We harnessed recent advances in bioacoustic technology to create a survey framework that integrates passive acoustic surveys and semi-automated detection to increase monitoring capacity for the orange-bellied parrot. We developed a custom BirdNET classifier for the orange-bellied parrot and compared efficacy of acoustic and field surveys using an occupancy framework. We deployed autonomous recording units across the orange-bellied parrot’s breeding range in southwest Tasmania and concurrently undertook between three and six repeated point-count surveys at the same 48 sites using human observers. Our custom BirdNET classifier had high accuracy and discrimination abilities. Validation of model scores across a week (5,712 hours of audio) required 60 hours reviewing time and yielded a 95% confidence of a correct BirdNET prediction at scores over 0.998. Occupancy analysis showed that the detection probability of acoustic surveys (*p* = 0.80) was more than eleven times greater than field surveys by skilled ecologists familiar with the species (*p* = 0.07). We provide a template for how to implement monitoring of the orange-bellied parrot and recommendations for how our methods can be improved to optimise the classifier to account for other species and locations.

## INTRODUCTION

Rare species are often characterised by major data deficiencies, which limit the ability to monitor populations, detect trends, and identify the threats driving declines. These challenges are compounded when species occur in remote or rugged landscapes (Wood et al. 2023), or are highly mobile and use sites only briefly during their life history (Runge et al. 2014). Under such conditions, tracking habitat use and dynamic ranges becomes difficult, making it harder to implement effective conservation actions (e.g., habitat protection, invasive species control). Improved survey methods can help overcome these limitations (Runge and Tulloch 2018).

Passive acoustic monitoring involves deploying autonomous recording units (ARUs) across landscapes to collect environmental audio recordings and species vocalisations (Sugai et al. 2019; Kahl et al. 2021; Teixeira et al. 2024). Detection algorithms can be subsequently applied to these recordings to identify distinctive vocalising species, enabling large-scale occupancy monitoring and ecological inference while minimizing disturbance (Wood et al. 2021). In conjunction with species-specific tools to enable efficient processing of large datasets, passive acoustic monitoring has a scalable capacity to identify range changes (Wood et al. 2019; Wood et al. 2024), breeding success (Teixeira et al. 2022), invasive species (Hofstadter et al. 2022; Bota et al. 2024), help direct observer-based surveys (Kramer et al. 2024) and guide rapid conservation interventions (Wood et al. 2024).

The orange-bellied parrot *Neophema chrysogaster* is a migratory, mobile species and one of Australia’s most critically endangered birds (Birdlife International 2018; DCCEEW Department of Climate Change 2026). The population has been intensely managed for over four decades but numerous uncertainties remain about the species’ movements, potential dispersal within its breeding range, and precise overwintering locations (Bussolini et al. 2026; Orange-bellied Parrot Tasmanian Program 2024). Occurrence surveys of orange-bellied parrots across their range have relied primarily on targeted field surveys, focused at sites where the species has been observed either recently or historically (DELWP 2016). However, limited resourcing means that spatiotemporal coverage of field survey efforts are restricted to a few accessible locations, sometimes resulting in long intervals between survey repeats.

Passive acoustic surveys may overcome some of the challenges and limitations of field survey approaches for this critically endangered bird. The increasing availability of ARUs that can record audio for weeks or months combined with advances in audio data processing (Kahl et al. 2021) have the potential to increase spatiotemporal survey coverage for the orange-bellied parrot. However, increased survey coverage is only beneficial if the resulting audio can be processed efficiently and accurately. A key data limitation for the orange-bellied parrot is the scarcity of labelled vocalisation recordings in public archives. Consequently this species is not included as a pre-trained class in BirdNET (Kahl et al. 2021), one of the most widely-used machine learning tools for bird sound identification. Fortunately, advances in transfer learning, the application of an algorithm to a task for which it was not explicitly designed, has proven quite successful in bioacoustics (Ghani et al. 2023). Without any machine learning expertise required, BirdNET can now be retrained to identify novel sounds, a potentially significant advance for data-deficient species that are generally lacking in training data. There is potential for dual advances in bioacoustics, increasing availability of ARUs and the ability to make custom classifiers, to (i) provide a passive acoustic monitoring workflow that can augment existing field survey efforts, and (ii) potentially increase the capacity to detect species presence at a site while reducing active field effort (Digby et al. 2013; Darras et al. 2018).

The aim of this study was to develop a passive acoustic survey method for the orange-bellied parrot that can both augment existing monitoring efforts and meet or exceed field observer-based surveying capacity. Our objectives were to: (i) use BirdNET to develop a custom orange-bellied parrot detector and (ii) evaluate how well passive acoustic monitoring can be used to detect occupied sites. To achieve this, we implemented an occupancy study in the breeding area at Melaleuca, southwest Tasmania using both field and passive acoustic monitoring-based survey methods. These outcomes could offer an efficient new approach for monitoring one of Australia’s most endangered birds across remote and logistically challenging habitats, thereby addressing important objectives for the species long-term recovery.

## METHODS

### Study species and area

The orange-bellied parrot is a small (45 g) migratory bird that breeds in southwest Tasmania during the austral spring/summer and overwinters on the coastal south-eastern Australian mainland (Bussolini et al. 2026). During the non-breeding season, birds occupy a range of habitats including coastal saltmarsh, and wetlands, and occupies buttongrass moorlands, melaleuca shrubland, as well as *Eucalyptus nitida* forest patches during the breeding season (Ehmke 2009; White et al. 2016; Brown and Wilson 1980). Orange-bellied parrots feed on a range of ground- and shrub-layer food plants (Brown and Wilson 1980; OBPRT 1999). Since 1988, supplementary food has been provided in the breeding area (Garnett, Szabo, and Dutson 2010; OBPRT 1999), and the majority of the population aggregates at feed tables during the breeding season (Riley 2025; Stojanovic et al. 2020).

We focussed on the breeding area for the orange-bellied parrot in the Melaleuca Valley, Southwest National Park, Tasmania, where the entire known wild population of orange-bellied parrots congregate. The landscape is dominated by buttongrass moorland with peatlands, sedgelands, and patches of temperate rainforest. We conducted the study during the orange-bellied parrot breeding season in Tasmania (January-February) when parrots were provisioning chicks or fledglings. We selected 48 survey sites across the study area with ≥400m separating survey points to ensure spatial independence.

### Field surveys

At each site, observers conducted repeated 10-minute occupancy surveys to record detection and non-detection data for orange-bellied parrots. An initial pilot trial indicated that 20-minute surveys did not improve naïve occupancy estimates. To avoid violating the assumption of site closure (MacKenzie 2006) we conducted three to six surveys per site (MacKenzie and Royle 2005) within a month from 29^th^ January 2025. We considered a site occupied if an orange-bellied parrot was detected, inclusive of birds flying through the site, because the species is readily detected by its distinctive flight call even if individuals are not sighted. Four skilled observers conducted surveys at all times of day but not in conditions likely to reduce detectability (e.g., rain, winds > 30km/hour). We did not apply a maximum distance cut-off to the observations.

### Passive acoustic monitoring

We deployed 48 ARUs (Frontier Labs, model: BAR-LT) at each site on the first visit and retrieved them on the last visit. Autonomous recording units were programmed to record continuously between one-hour before sunrise to one-hour after sunset via a single omni-directional microphone, pointing toward the ground to reduce interference with wind. Units were set to a sample rate of 48 kHz, 16-bit resolution and gain of +40 dB. Recordings were a mean of 24.3 days per unit (range: 19– 32 days). All units recorded simultaneously for a 19-day period from 1^st^-19^th^ February 2025. In total, we recorded 19,565 hours of audio across all sites.

### Analyses

We developed a custom classifier using the BirdNET algorithm v2.4 (Kahl et al. 2021), a convolutional neural network trained on sound libraries to identify species based on 3-second audio segments. Ghani et al. (2023) showed that BirdNET v2.4 can be used in a transfer learning context to develop custom classifiers, a valuable extension of BirdNET particularly for species like the orange-bellied parrot that are data-deficient and not included as a pre-trained class. We compiled training recordings from passive acoustic surveys. The training dataset comprised 19 hours of recordings, including 15 one-hour files from field deployments and four one-hour files from captive recordings.

We manually reviewed and annotated recordings to identify orange-bellied parrot vocalisations suitable for training (i.e., high signal-to-noise ratio) using Raven Pro v1.6 (K. Lisa Yang Center for Conservation Bioacoustics at the Cornell Lab of Ornithology 2026). Training data were iteratively curated to improve classifier performance. For example, to improve model performance, we removed calls that overlapped strongly with other species. This yielded 553 annotated orange-bellied parrot training selections. We created additional non-target classes for species that caused frequent false positives during preliminary analyses (including grey fantail *Rhipidura albiscapa*, superb fairywren *Malurus cyaneus*, tree martin *Petrochelidon nigricans*, crescent honeyeater *Phylidonyris pyrrhopterus*, and New Holland honeyeater *Phylidonyris novaehollandiae*) and included these as additional classes in our custom model to improve discrimination among acoustically similar signals. Environmental sounds (e.g., rain, insects, wind, and other background noises) were collated into a single noise class. We included multiple vocalisation types of the orange-bellied parrot in the training data, inclusive of the flight call, chatter/buzz calls and whistle (Figure 1).

**Figure 1.**
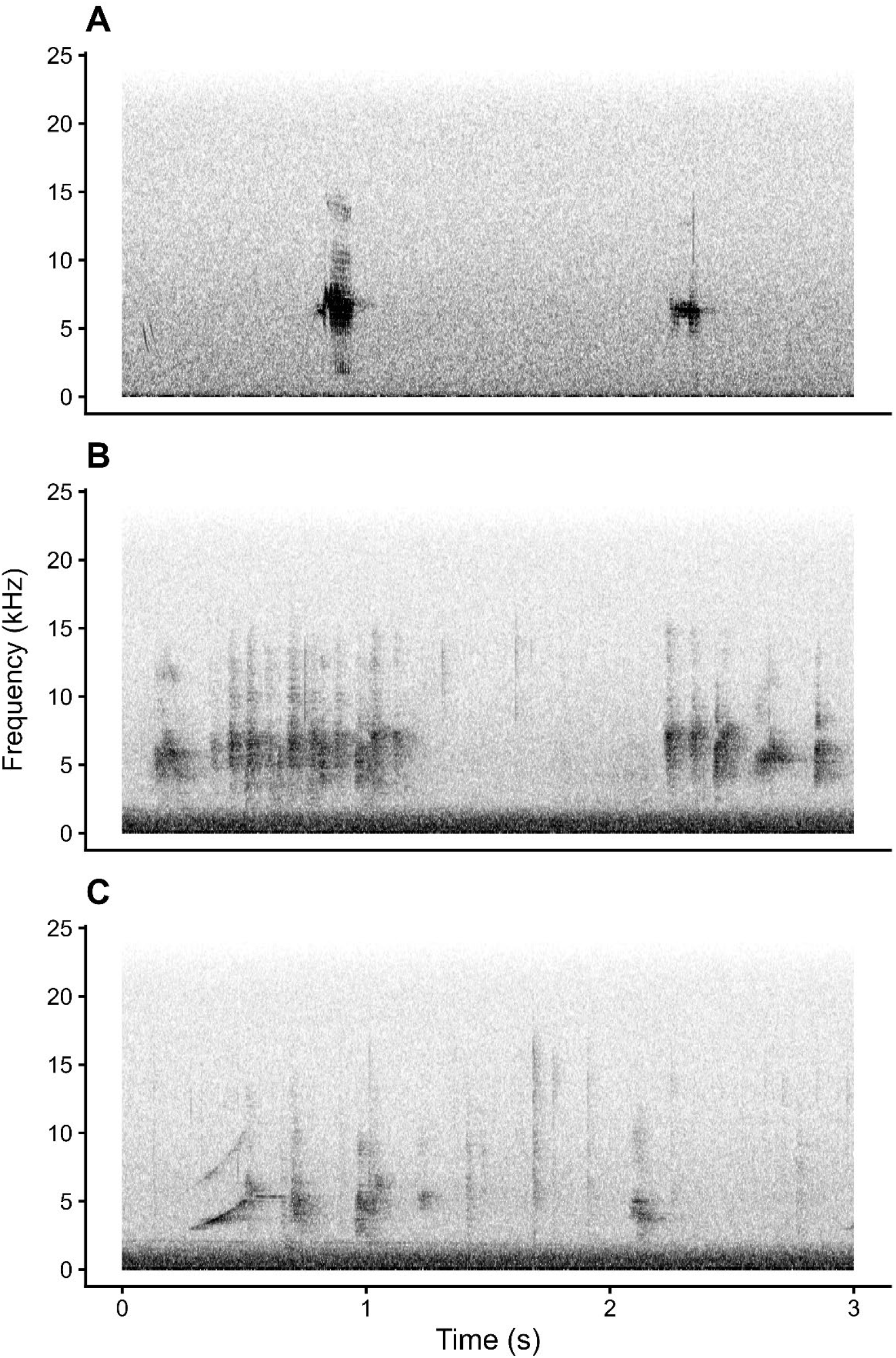
Examples of Orange bellied parrot vocalisations used to train the custom BirdNET classifier. (a) flight call, b) chatter (buzz call), and whistle.

We evaluated the classifier using a separate test dataset comprising seven one-hour recordings from field deployments in Melaleuca. To prevent data leakage that might bias the evaluation process, our test data and training data were obtained from different dates and times. We analysed the test data with our custom version of BirdNET and extracted predictions (i.e., three-second audio segments).

We conducted segment extraction twice to capture both the full score distribution (confidence 0.1– 1.0) and high-confidence predictions (confidence 0.9–1.0) (total segments = 200). We manually reviewed any segments predicted to contain an orange-bellied parrot to determine whether each represented a true or false positive. These validated predictions were subsequently used to evaluate the classifier performance. Evaluation metrics included the area under the receiver operating characteristic curve (AUROC) and average precision (AP), representing the area under the precision– recall curve. We further evaluated the pilot classifier performance on the test dataset using logistic regression per Wood and Kahl (2024). We next processed the full 19,565-hour acoustic dataset with our custom classifier. To calibrate a review threshold for the classifier suitable for large datasets, validation was required to confirm orange-bellied parrot vocalisations by manually listening to recordings. We applied a standardized reviewing process and manually reviewed the ten highest-scoring BirdNET predictions per site and day for a seven-day subset of the data where all autonomous recording units were operational (5,712 h of recordings). BirdNET produces unitless “confidence scores” that reflects how confident the algorithm is in a given prediction (Wood and Kahl 2024). Higher scores generally indicate a greater likelihood that the prediction is correct, but the relationship between score and prediction accuracy varies among species, recording conditions, and even among studies of the same species.

For each prediction we scored the orange-bellied parrot outcome as a binomial value for true or false (1/0) and if 0, we categorised false positives (i.e., wind, insect, bird, noise) and identified non-target bird species. We analysed results from the acoustic validation in R (R Core Team 2025) and evaluated the classifier performance on the field dataset using logistic regression per Wood and Kahl (2024). Specifically, confidence scores were transformed using the logit function:

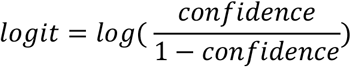

We then fitted three binomial generalized linear models per BirdNET score use recommendations of Wood & Kahl (2024).

1. A null model including only an intercept,
2. A model using the confidence score as predictor, and
3. A model using the back-transformed logit score.

We compared model performance using Akaike’s Information Criterion (AIC). The top-ranked model was then used to estimate the probability that a BirdNET prediction represented a correct prediction (i.e., a detection of our target species) and to calculate threshold confidence values corresponding to probabilities of 0.90, 0.95 and 0.99.To evaluate the efficacy of passive acoustic surveys relative to field surveys, we conducted occupancy modelling using R package ‘unmarked’ v.1.5.1 (Kellner et al. 2023; Fiske and Chandler 2011) following the framework of MacKenzie et al. (2002). Data from both the field surveys and the validated acoustic survey data were included. We included a binomial presence value of each survey occasion. For the field surveys, a single survey occasion was defined as a ten-minute survey. A single survey occasion in the acoustic data was included, which collapsed daily acoustic data into a single presence/absence value for each day.

We fit a single-season occupancy model with 13 survey occasions (6 human observer, 7 acoustic). For detection probability (*p*) we fitted both a null model (i.e., *p* was uniform) and a model including the survey method (field surveys vs. passive acoustic monitoring) as a fixed effect. We compared models based on ΔAIC. We did not include date because this variable was confounded in the acoustic data by a short time window (one week), whereas the field data took place over 3-4 weeks. Likewise, given the continuous recording of the acoustic data, we could not include a time for absence data, thus, we excluded time from the occupancy analysis.

This study was approved by the [redacted for review] Ethics Committee (ethics approval number redacted for review), with permit number [redacted for review]. All ARU deployment locations were selected away from visitor infrastructure.

## RESULTS

### Objective 1: Development of a custom classifier

The BirdNET-based orange-bellied parrot classifier demonstrated strong discrimination between classes in our independent test data. The receiver operating characteristic analysis showed a high area under the curve (AUROC = 0.908), indicating that the classifier effectively distinguished between true positive and false positive observations across a range of confidence thresholds (Figure 2a). Similarly, the precision–recall evaluation produced a high average precision score (AP = 0.949), demonstrating that the model maintained high precision (proportion of predictions that are correct) while retaining strong recall (proportion of target signals present and correctly identified) (Figure 2b). However, as expected, precision decreased at higher recall values (i.e., more false positives occur with higher recall at lower score thresholds).

**Figure 2.**
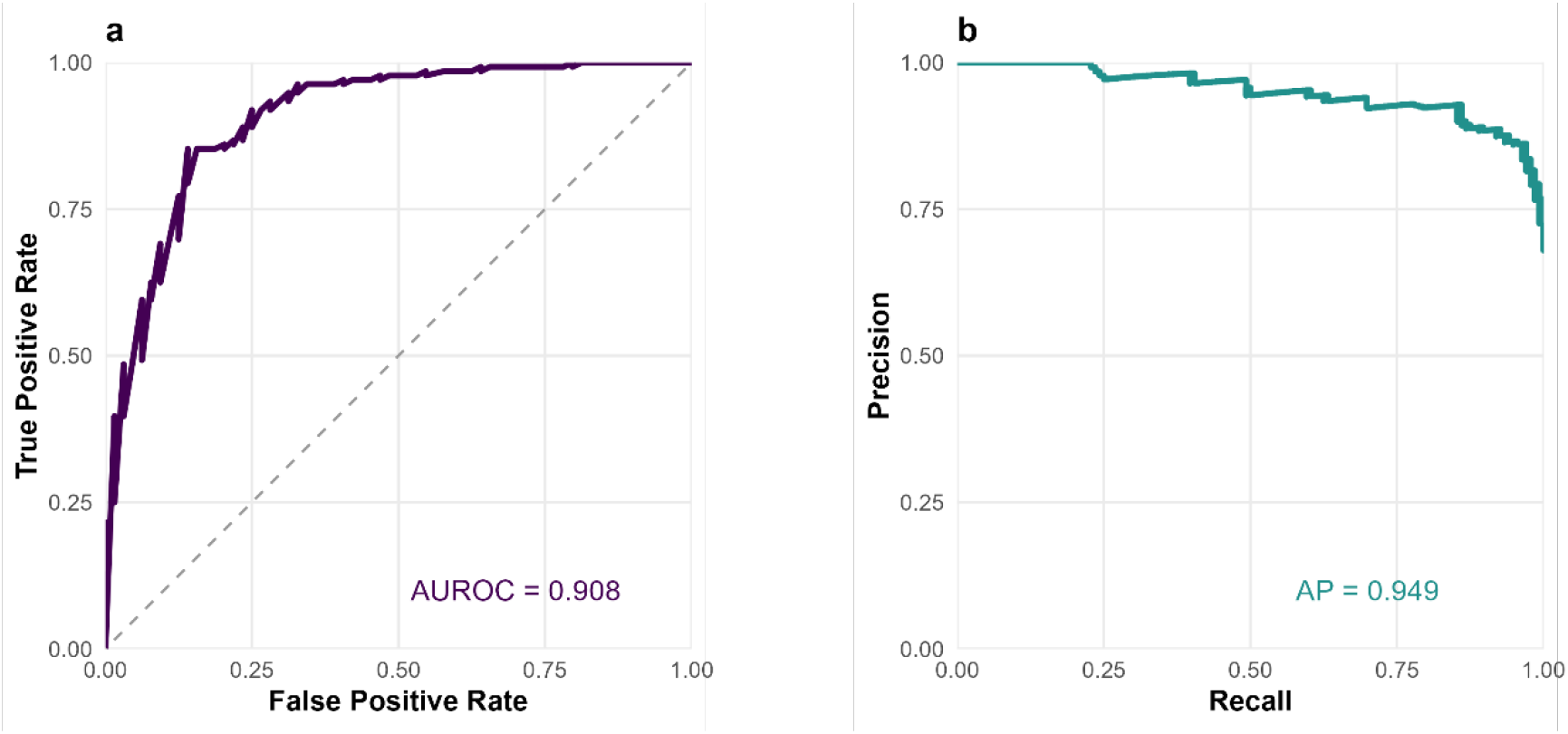
Performance of the BirdNET classifier for the orange-bellied parrot. (a) Receiver operating characteristic curve showing true positive vs. false positive rate trade-off across confidence thresholds. The area under the receiver operating curve (AUROC) summarises classifier performance across all thresholds. The dashed line indicates the performance of a random classifier. (b) Precision–recall curve showing the relationship between precision and recall across confidence thresholds. Average precision (AP) summarises the ability to detect the species while minimising false detections.

**Figure 3.**
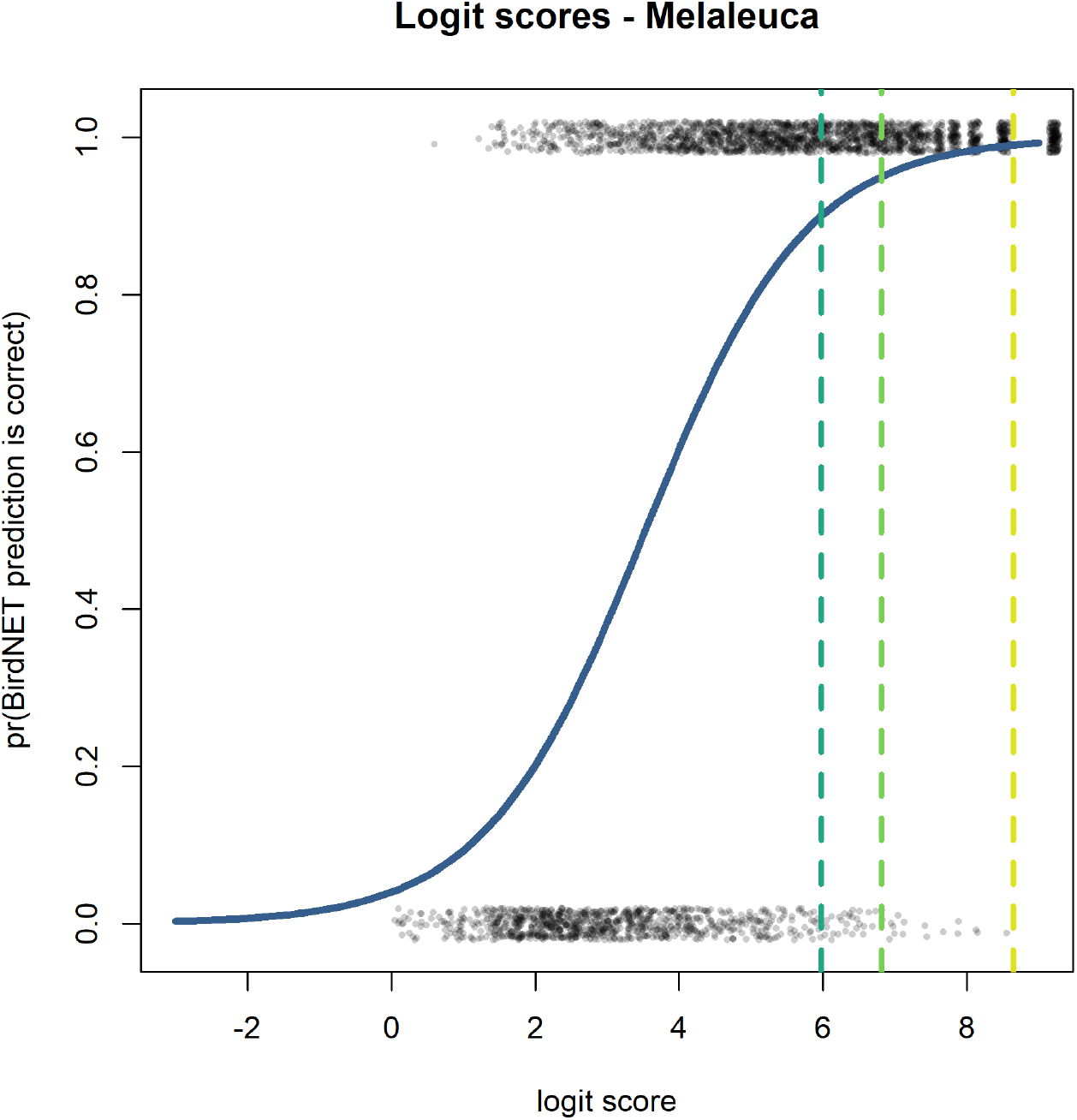
Logistic regression estimates of correct BirdNET predictions (blue line) for validated predictions (grey points) on the logit scale (preferred model) for the orange-bellied parrot. Confidence thresholds for the probability of a true positive prediction are shown in dashed lines for 0.9% (cyan), 0.95% (green) and 0.99% (yellow) accuracy.

Logistic regression based on data from the seven-day validated dataset confirmed that high classifier confidence scores indicated correct predictions. AIC-based model comparison favoured the logit-transformed model over the confidence score model and the null model (Table 1). Reverse transformation of logit thresholds to confidence scores indicated that a predicted probability of 0.95 of a true positive corresponded to a confidence score 0.9989 (Table 2).

**Table 1.** AIC ranking of logistic models predicting correct orange-bellied parrot call identification from BirdNET confidence scores in the 1-week validated Melaleuca field dataset (comprising 5,712 hours audio).

| Model | AIC | Delta AIC |
| --- | --- | --- |
| Logit model | 2393.25 | 0.00 |
| Confidence model | 2723.81 | 330.56 |
| Null model | 3717.32 | 1324.07 |

**Table 2.** Thresholds of the probability of a true positive (TP) and BirdNET logit and confidence score that yields the given probability for the orange-bellied parrot in the Melaleuca Valley, Southwest Tasmania. Confidence scores for each logit were obtained by back-transforming.

| pr(TP) | Logit | Confidence Score |
| --- | --- | --- |
| 0.90 | 5.98 | 0.9974 |
| 0.95 | 6.81 | 0.9989 |
| 0.99 | 8.65 | 0.9998 |

Validating the ten highest-scoring predictions per site per day for one week of audio data required analyst time of an average 82 minutes per site and 60 hours across all 48 sites.

### Objective 2: Passive acoustic detection of occupied sites

Naïve occupancy was three times higher with acoustic surveys (0.94) than field surveys (0.29). Orange-bellied parrot vocalisations were detected via passive acoustic monitoring at 45/48 sites in the one week of validate audio data we processed with the custom classifier (Figure 4) (*n* detections = 251, *n* acoustic survey days = 336). In contrast, field surveys detected orange-bellied parrots at just 14/48 sites (*n* detections = 28, *n* field surveys = 184).

**Figure 4.**
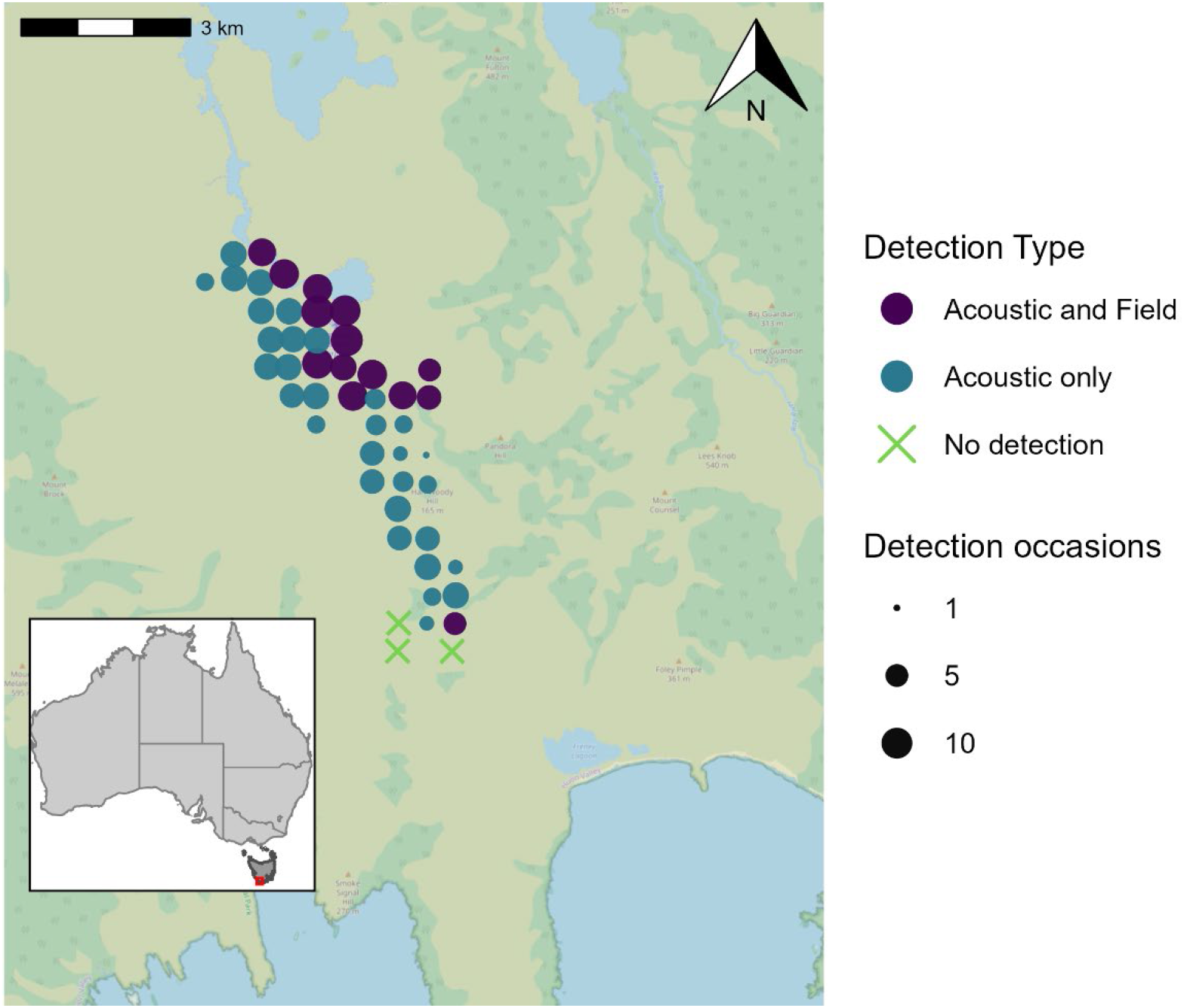
Map of the study area in the Melaleuca Valley, showing the number and type of orange-bellied parrot detections derived from passive acoustic monitoring and observer-based field surveys. The study area is in south-western Tasmania, Australia.

All sites with a field survey-derived detection were corroborated with at least one passive acoustic survey detection (Figure 4). The majority of sites with a field survey derived detection (13/14) were close to the location of supplementary feeding stations. Passive acoustic monitoring detected orange-bellied parrots across a much broader extent of the study area.

In the occupancy analysis, model selection strongly supported inclusion of survey method (field surveys vs. passive acoustic monitoring) as a detection covariate (w = 1.0; ΔAIC null model = 195.91). Detection probability via passive acoustic monitoring (0.80 ± 0.07) was 11 times that of field surveys (0.07 ± 0.03) (Table 3).

**Table 3:**
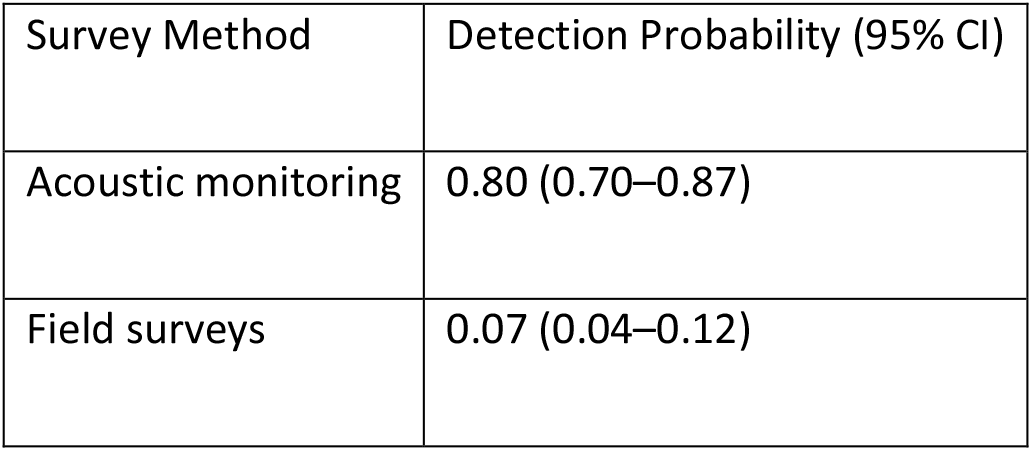
Survey Method Detection Probability (95% CI)

The cumulative, or seasonal, detection probability (*p\**) across the one-week acoustic survey period was >0.99, meaning our approach yielded near-certain detection of orange-bellied parrots if present. It would take approximately 64 observer-based surveys to achieve a similar level of confidence in site occupancy. Overall occupancy probability (Ψ) was 0.94, which matched naïve occupancy from acoustic survey data.

## DISCUSSION

Detecting rare species in the field is inherently challenging, so finding effective survey methods can be the difference between extinction and conservation success for highly imperilled species. In this study we assessed the efficacy of acoustic monitoring for the Critically Endangered orange-bellied parrot, a species for which surveys by human observers are known to yield only limited and patchy data in field conditions (Stojanovic et al. 2017). Using an occupancy modelling framework applied within the species’ core breeding area, we demonstrate that our passive acoustic monitoring approach, powered by a custom BirdNET classifier, dramatically outperformed field surveys by skilled observers.

We confirmed presence of orange-bellied parrots at the majority of sites using passive acoustic monitoring, with a 0.95 probability of a true positive at a confidence score that approached ≥0.9989. Passive acoustic monitoring yielded a detection probability of 0.8 for a one-day deployment, which was more than 11-fold greater than comparable surveys by skilled field ecologists over the same time period. This result is an important step toward developing a new approach to monitoring the orange-bellied parrot and shows that detecting this species in field conditions is possible using the approach we have developed.

Our classifier for the orange-bellied parrot had very high performance, effectively distinguishing between positive and negative observations across a range of BirdNET confidence score thresholds indicated by the AUROC. Average precision was also very high. Predictions within the high-confidence segments were mostly correct, with more false positives observed at lower confidence scores. The trade-off in increasing the confidence score threshold was sacrificing recall, as high thresholds filter out lower scoring true-positive orange-bellied parrot calls. In our case, the objective of the survey was presence-data, and it was preferrable to prioritise precision over recall, as if the site is occupied, there will likely be high scoring vocalisations in the same dataset as low scoring vocalisations.

The classifier performed well when applied to the one-week survey dataset, though a slightly higher confidence threshold was needed to achieve 95% precision compared to the pilot (0.9989 vs. 0.987). This likely reflects greater false positives in the field data, which spanned more locations, conditions, and non-target species diversity. These findings align with previous work showing that automated classifiers can aid detection in natural soundscapes but performance varies across environments and dataset sizes (Sossover et al. 2024; Teixeira et al. 2022; Sugai et al. 2019) and is often limited by background noise, overlapping species calls, and variation in call quality (Pérez Granados 2023; Priyadarshani, Marsland, and Castro 2018).

Passive acoustic monitoring has emerged as a popular approach to monitoring cryptic species, but this method is not necessarily a panacea. From the perspective of conservation of the orange-bellied parrot, managers must evaluate the costs and benefits of alternative methods, which our study can now clarify. To illustrate this, consider a scenario where a land manager is seeking to deliver an occupancy-style survey of orange-bellied parrots in an area of potential habitat. Field surveys by skilled ecologists have the advantage of being relatively low-cost in terms of delivery timelines; an ecologist might visit sites 3-5 times to implement short surveys, then analysis of the data can be conducted quickly without need for post-processing of field data. However, our results show that the probability of even a very skilled observer detecting orange-bellied parrots in a relatively short survey window is very low, which greatly increases the risk of false negatives. In contrast, passive acoustic monitoring has higher up-front costs than field surveys. This is attributable to both the cost of purchasing recorders and personnel costs to deploy and retrieve recorders. While these costs are comparable to two site visits, passive acoustic monitoring also needs to budget for the desktop validation of sound files. However, once these up-front costs are accounted for, passive acoustic monitoring can dramatically improve detection probability of orange-bellied parrots while reducing the need for repeated site visits.

Given that the orange-bellied parrot population is unable to absorb any additional mortality from the emergence of new threats across their range (Stojanovic and Bussolini 2026), the risk of failure to detect the species when present may be severe. The reduction of false negatives afforded by passive acoustic monitoring may make the higher upfront costs a worthwhile investment. Our results indicate that a one-week acoustic deployment yielded a cumulative detection probability exceeding 0.99, while achieving a similar level of certainly using observer-based surveys would require approximately 64 independent site visits. This difference is particularly relevant for applications such as development impact assessment, habitat evaluation, and recovery planning, where failure to detect the species when present may lead to inappropriate management decisions. Although there were strong results in this study, several points deserve consideration. Our study was conducted within the core orange-bellied parrot breeding range at Melaleuca. While our iterative training of the pilot classifier yielded strong performance on this dataset, its performance may vary in other regions. In this study area, the orange-bellied parrot is the only *Neophema* present, but in other parts of their range – during migration and wintering on the mainland – multiple species from this genus may co-occur. This increases the potential for misclassification as congeners produce acoustically similar calls. Further work is needed to develop custom call classifiers incorporating vocalisations from other *Neophema* species, as well as those of non-target species that occur across the full migratory range and pose additional risks of false positives.

Importantly, false negatives cannot be completely ruled out using acoustic monitoring. This risk is only reduced relative to relying solely on observed-based field surveys. Multiple factors may contribute to false absences when a species is present while using ARUs, inclusive of behavioural factors, environmental noise, limited recorder range, equipment failure, or unsuitable sampling design. Nevertheless, acoustic monitoring is a powerful tool that can be used to direct limited field effort toward sites where follow-up surveys are most needed.

Developing effective techniques to detect and monitor rare species is a fundamental conservation need. Our study demonstrates that passive acoustic monitoring combined with transfer learning using BirdNET can significantly outperform traditional observer-based surveys for detecting the Critically Endangered orange-bellied parrot. It provides an example of how careful development and comparison of survey methods can support the implementation of species’ recovery actions by increasing detection probability.

## Ethics statement

We conducted research under [redacted for review] animal ethics license (redacted for review), Tasmanian Scientific Permit (redacted for review), Tasmanian Parks and Wildlife approvals, Aboriginal Heritage and Cultural Heritage approval.

## Acknowledgements

We would like to acknowledge the following organisations and groups for their support in delivering this project: Orange-bellied Parrot Tasmanian Program, Department of Natural Resources and Environment (NRE) Tasmania, Orange-bellied Parrot Recovery Team, Tasmanian Parks and Wildlife, Zoos Victoria, Department of Energy, Environment and Climate Action (DEECA), Par Avion, and Frontier Labs Australia. Thank you to the following individuals: Shannon Troy, Clare Lawrence, Madalyn Riley, Jennifer Mudge, Michael Domrose, Lisa McKay, Nicole Gill, Marike Oliphant, Darcy Whittaker, Jessica Willemse, Michael Maggs, Kieran Aland, Otis Mayocchi, Michael Magrath, Sarah Agterhuis, Kerri Duncan, and Rachel Pritchard. Funding was provided by the Department of Climate Change, Energy, the Environment and Water ‘Saving our Native Species’ Program. The authors have no conflicts of interest to declare.

## Author Contributions

Conceptualisation: DS, GO, CW; Methodology: GO, CW, DS; Data curation: GO, TH, LB, AK; Investigation: all authors; Validation: GO, TH; Formal analysis: GO; Project administration: DS, GO; Resourcing: DS; Funding acquisition: DS, FA; Writing – original draft: GO, DS, CW; Writing – review and editing: all authors.

## References

Birdlife International 2018, ‘Neophema chrysogaster, In The IUCN Red List of Threatened Species 2018: eT22685203A130894893.’.

Bota, G, Manzano-Rubio, R, Fanlo, H, Franch, N, Brotons, L, Villero, D, Devisscher, S, Pavesi, A, Cavaletti, E & Pérez-Granados, C 2024, ‘Passive acoustic monitoring and automated detection of the American bullfrog’, Biological Invasions, vol. 26, no. 4, pp. 1269–79.

Brown, P. & Wilson, RI 1980, ‘A survey of the Orange-bellied Parrot (Neophema chrysogaster) in Tasmania, Victoria, and South Australia. Report prepared for World Wildlife Fund (Australia). National Parks and Wildlife Service, Tasmania, Australia.’.

Bussolini, LT, D. Gautschi, G. V. Hodder, G. Owens, S. Troy & Stojanovic, D 2026, ‘Orange-bellied Parrot (Neophema chrysogaster), version 2.0. In Birds of the World (M. G. Smith and G. M. Kirwan, Editors).’, Cornell Lab of Ornithology, Ithaca, NY, USA.

Darras, K, Batáry, P, Furnas, B, Celis-Murillo, A, Van Wilgenburg, SL, Mulyani, Y. & Tscharntke, T 2018, ‘Comparing the sampling performance of sound recorders versus point counts in bird surveys: A meta-analysis’, Journal of Applied Ecology, vol. 55, no. 6, pp. 2575–86.

DCCEEW Department of Climate Change, E, the Environment, and Water 2026, Neophema chrysogaster-Orange-bellied parrot, In Species Profile and Threats Database.

DELWP Department of Environment Land Water and Planning 2016, National Recovery Plan for the Orange-bellied Parrot, Neophema chrysogaster, Australian Government, Canberra, Australia.

Digby, A, Towsey, M, Bell, B. & Teal, PD 2013, ‘A practical comparison of manual and autonomous methods for acoustic monitoring’, Methods in Ecology and Evolution, vol. 4, no. 7, pp. 675–83.

Ehmke, G 2009, ‘Potential occurrence and optimal habitat models for the Orange-bellied parrot in south-eastern mainland Australia. Birds Australia, Melbourne, Australia.’.

Fiske, I & Chandler, R 2011, ‘unmarked: An R package for fitting hierarchical models of wildlife occurrence and abundance’, Journal of statistical software, vol. 43, no. 10.

Garnett, JT, Szabo, J. & Dutson, G 2010, The Action Plan for Australian Birds 2010, CSIRO Publishing, Melbourne, Australia.

Ghani, B, Denton, T, Kahl, S & Klinck, H 2023, ‘Global birdsong embeddings enable superior transfer learning for bioacoustic classification’, Scientific Reports, vol. 13, no. 1, p. 22876.

Hofstadter, DF, Kryshak, NF, Wood, CM, Dotters, BP, Roberts, KN, Kelly, KG, Keane, JJ, Sawyer, SC, Shaklee, PA, Kramer, HA, Gutiérrez, R. & Peery, MZ 2022, ‘Arresting the spread of invasive species in continental systems’, Frontiers in Ecology and the Environment, vol. 20, no. 5, pp. 278–84.

K. Lisa Yang Center for Conservation Bioacoustics at the Cornell Lab of Ornithology 2026, ‘Raven Pro: Interactive Sound Analysis Software (Version 1.6.5) [Computer software].’, Ithaca, NY: The Cornell Lab of Ornithology.

Kahl, S, Wood, CM, Eibl, M & Klinck, H 2021, ‘BirdNET: A deep learning solution for avian diversity monitoring’, Ecological Informatics, vol. 61, p. 101236.

Kellner, KF, Smith, AD, Royle, JA, Kéry, M, Belant, J. & Chandler, RB 2023, ‘The unmarked R package: Twelve years of advances in occurrence and abundance modelling in ecology’, Methods in Ecology and Evolution, vol. 14, no. 6, pp. 1408–15.

Kramer, HA, Kelly, KG, Whitmore, SA, Berigan, WJ, Reid, DS, Wood, CM, Klinck, H, Kahl, S, Manley, PN, Sawyer, S. & Peery, MZ 2024, ‘Using bioacoustics to enhance the efficiency of spotted owl surveys and facilitate forest restoration’, The Journal of Wildlife Management, vol. 88, no. 2, p. e22533.

MacKenzie, DI 2006, Occupancy estimation and modeling : inferring patterns and dynamics of species occurrence, 1 edn., Elsevier, Amsterdam; Boston.

MacKenzie, DI, Nichols, JD, Lachman, GB, Droege, S, Andrew Royle, J & Langtimm, CA 2002, ‘Estimating site occupancy rates when detection probabilities are less than one’, Ecology (Durham), vol. 83, no. 8, pp. 2248–55.

MacKenzie, D. & Royle, JA 2005, ‘Designing occupancy studies: general advice and allocating survey effort’, The Journal of Applied Ecology, vol. 42, no. 6, pp. 1105–14.

OBPRT Orange-bellied Parrot Recovery Team 1999, Orange-bellied Parrot Recovery Plan 1998-2002., Department of Primary Industries and Water (DPIW), Hobart.

Orange-bellied Parrot Tasmanian Program 2024, Orange-bellied Parrot Migration Tracking-Interim Report 2024.

Pérez Granados, C 2023, ‘A First Assessment of Birdnet Performance at Varying Distances: A Playback Experiment’, Ardeola, vol. 70, pp. 257–69.

Priyadarshani, N, Marsland, S & Castro, I 2018, ‘Automated birdsong recognition in complex acoustic environments: a review’, Journal of Avian Biology, vol. 49.

R Core Team 2025, ‘R: A Language and Environment for Statistical Computing’, R Foundation for Statistical Computing.

Riley, M, Lawrence, C., Troy, S. 2025, Report on the Melaleuca wild population 2024/25, (Orange-bellied Parrot Tasmanian Program, chairman), Department of Natural Resources and Environment Tasmania, Hobart, Tasmania.

Runge, C & Tulloch, AIT 2018, ‘Solving problems of conservation inadequacy for nomadic birds’, Australian Zoologist, vol. 39, no. 2, pp. 280–95.

Runge, CA, Martin, TG, Possingham, HP, Willis, S. & Fuller, RA 2014, ‘Conserving mobile species’, Frontiers in Ecology and the Environment, vol. 12, no. 7, pp. 395–402.

Sossover, D, Burrows, K, Kahl, S & Wood, CM 2024, ‘Using the BirdNET algorithm to identify wolves, coyotes, and potentially their interactions in a large audio dataset’, Mammal Research, vol. 69, no. 1, pp. 159–65.

Stojanovic, D, Alves, F, Cook, H, Crates, R, Heinsohn, R, Peters, A, Rayner, L, Troy, S. & Webb, MH 2017, ‘Further knowledge and urgent action required to save Orange-bellied Parrots from extinction.’, Emu-Austral Ornithology vol. 18, pp. 126–34.

Stojanovic, D & Bussolini, LT 2026, ‘How Many Deaths Are Too Many? Assessing the Impact of Additional Mortality on a Critically Endangered Bird’, Animal Conservation, vol. n/a, no. n/a.

Stojanovic, D, Potts, J, Troy, S, Menkhorst, P, Loyn, R & Heinsohn, R 2020, ‘Spatial bias in implementation of recovery actions has not improved survival of Orange-bellied Parrots Neophema chrysogaster.’, Emu, vol. 120(3):263–268.

Sugai, LSM, Silva, TSF, Ribeiro, JW, Jr. & Llusia, D 2019, ‘Terrestrial Passive Acoustic Monitoring: Review and Perspectives’, BioScience, vol. 69, no. 1, pp. 15–25.

Teixeira, D, Linke, S, Hill, R, Maron, M & van Rensburg, BJ 2022, ‘Fledge or fail: Nest monitoring of endangered black-cockatoos using bioacoustics and open-source call recognition’, Ecological Informatics, vol. 69, p. 101656.

Teixeira, D, Roe, P, van Rensburg, BJ, Linke, S, McDonald, PG, Tucker, D & Fuller, S 2024, ‘Effective ecological monitoring using passive acoustic sensors: Recommendations for conservation practitioners’, Conservation Science and Practice, vol. 6, no. 6, p. e13132.

White, M, Menkhorst, P, Griffioen, P, Green, B, Salkin, O & Pritchard, R 2016, Orange-bellied Parrot: A retrospective analysis of winter habitat availability, 1985–2015, Arthur Rylah Institute for Environmental Research Technical Report Series Number 277, DELWP Department of Environment, Land, Water and Planning.

Wood, CM, Champion, J, Brown, C, Brommelsiek, W, Laredo, I, Rogers, R & Chaopricha, P 2023, ‘Challenges and opportunities for bioacoustics in the study of rare species in remote environments’, Conservation Science and Practice, vol. 5, no. 6, p. e12941.

Wood, C. & Kahl, S 2024, ‘Guidelines for appropriate use of BirdNET scores and other detector outputs’, Journal of Ornithology, vol. 165, no. 3, pp. 777–82.

Wood, CM, Klinck, H, Gustafson, M, Keane, JJ, Sawyer, SC, Gutiérrez, R. & Peery, MZ 2021, ‘Using the ecological significance of animal vocalizations to improve inference in acoustic monitoring programs’, Conservation Biology, vol. 35, no. 1, pp. 336–45.

Wood, CM, Popescu, VD, Klinck, H, Keane, JJ, Gutiérrez, RJ, Sawyer, S. & Peery, MZ 2019, ‘Detecting small changes in populations at landscape scales: a bioacoustic site-occupancy framework’, Ecological Indicators, vol. 98, pp. 492–507.

Wood, CM, Socolar, J, Kahl, S, Peery, MZ, Chaon, P, Kelly, K, Koch, RA, Sawyer, S. & Klinck, H 2024, ‘A scalable and transferable approach to combining emerging conservation technologies to identify biodiversity change after large disturbances’, Journal of Applied Ecology, vol. 61, no. 4, pp. 797–808.

